# Artificial Alzheimer’s – visualizing the process of memory decay, confusion and death in a simple Neural Network Model of Associative Memory

**DOI:** 10.1101/2022.05.20.492604

**Authors:** Lennart Gustafsson

## Abstract

This paper analyzes the behaviour of Alzheimer’s disease simulation in artificial neural networks and thereby suggests a new possible diagnosis for Alzheimer. Alzheimer’s disease is one of the most common diseases, increasing severity over time. Despite its high prevalence and thousands of yearly publications in this area, no cure has been found to date and early detection is one of the most important factor towards fighting the disease and possibly finding a cure. This paper analyses the behaviour of simulated Alzheimer in Hopfield memories and observes that the main factor influenced by the loss of connections is the time needed to recognize distorted symbols - while the recognition performance stays surprisingly high for a long time. Hence, I suggest a new early diagnosis approach which is based on slightly distorted elements (e.g., characters) and measures the time needed to perform the task, instead of just focusing on the subject’s recognition performance. This insight could have a large impact to future diagnose setups, once being validated in a clinical study.

## 1 Introduction

Alzheimer’s disease is characterized by cognitive deficiencies increasing over time due to spreading loss of neural tissue. There is, however, a degree of resilience in the functioning of the brain and it is well established that considerable damage can be done to the brain in an individual before Alzheimer’s disease is detected. There is presently no cure for Alzheimer’s disease but in hope that a cure will be found methods for detecting Alzheimer’s disease at an early stage is a field of research in its own right.

Alzheimer’s disease has been simulated [1] [2] using Hopfield memories, simple Artificial neural network models of associative memory [3]. If images have been stored in the memory matrix then a distorted image can be presented to the neural network and the stored image can be retrieved through a recurrent process in the neural network, assuming the distortion is not excessive.

Simulating Alzheimer’s disease consists of nulling elements in the memory matrix of the Hopfield memory, corresponding to eliminating synapses in a biological neural network, and presenting distorted images to this deteriorated neural network model.

## 2 Simulation Experiments and Results

### 2.1 Simulation set-up

In Figure 1 three rows of five objects are shown. The top row shows five logos created by the Swedish artist Tex Berg, each comprising three colored triangles. This row is stored in a Hopfield network. Each logo is described by a 153 element vector, determined from the positions of the triangles’ vertices and colors.

**Fig. 1.**
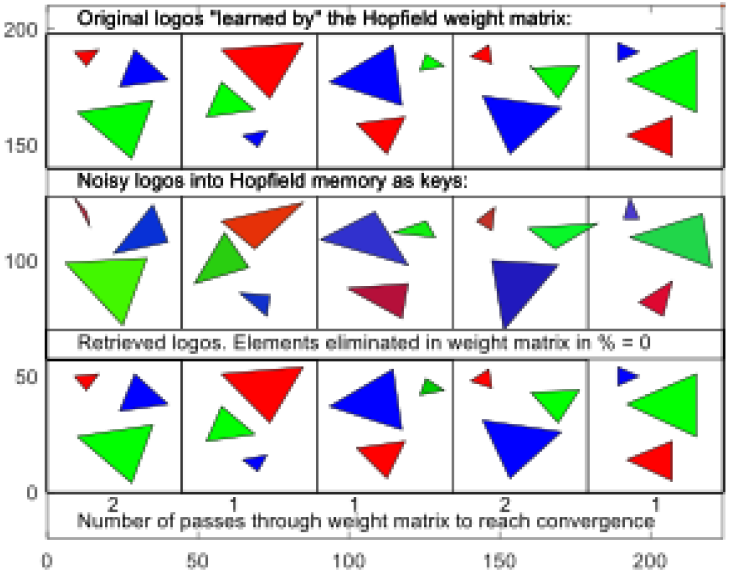
Memorized logos (top row), moderately distorted logos as inputs to Hopfield network (middle row) and retrieved logos (bottom row).

The middle row shows the same logos, moderately distorted, which are employed as inputs, keys, to the Hopfield network. The network should then through a recurrent process produce outputs identical to the stored logos.

In the bottom row this has successfully occurred. In this case no elements, or synapses, have been deleted. Since the keys are distorted logos they must be “corrected” and this is performed by passing the distorted input vector through the weight, or memory, matrix. One passage is sufficient for three of the keys, while for two cases an additional correcting loop through the weight matrix is also necessary.

In Figure 2 the inputs are even more distorted versions of the logos. In this case two of the logos, the first and the fourth, were not restored but “spurious attractors” were found. Convergence was reached in all cases and rather quickly at that (the number of passes through the weight matrix were not large). It is not surprising that convergence was reached since there is a theorem stating that a complete weight matrix guarantees convergence [3].

**Fig. 2.**
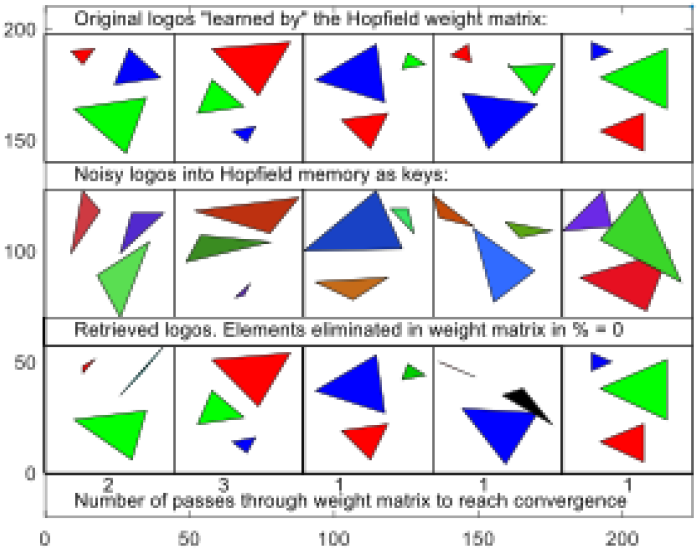
Memorized logos (top row), heavily distorted logos as inputs to Hopfield network (middle row) and resulting outputs after convergence (bottom row). No synapses were deleted.

### 2.2 The effect of deleting elements in the weight matrix i.e. synapses

In Figure 3 we show the effects when the inputs are as heavily distorted versions of the logos as in Figure 2 and synapses have been removed, in this case 50%.

**Fig. 3.**
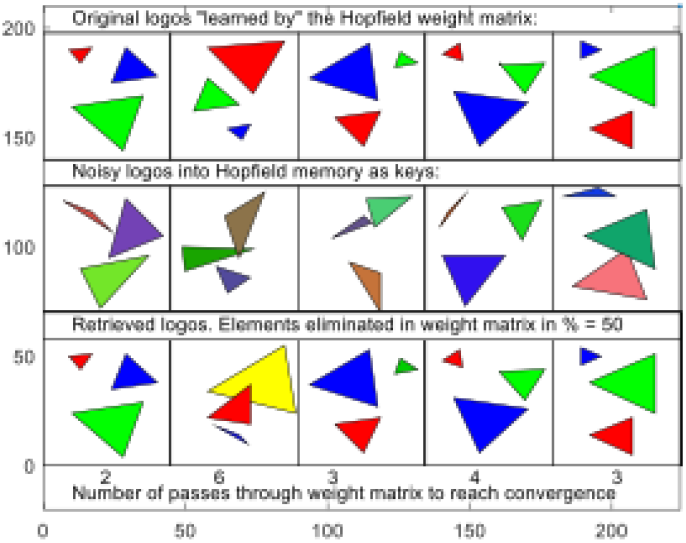
Memorized logos (top row), heavily distorted logos as inputs to Hopfield network (middle row) and resulting outputs after convergence (bottom row). 50 % of synapses were deleted.

Even though a substantial number of synapses have been removed in the case shown in Figure 3 the convergence results are not inferior to those in Figure 2. This is important – the quality of the results doesn’t deteriorate when synapses “start to be deleted”. It should be pointed out that the neural network in this case has a very large “overcapacity” – more than 20 logos could be accommodated in this neural network [4]. This means that the resilience against synapse loss is very great indeed.

The other important observation to be made from Figure 3 is that the convergence process is much longer than in Figure 2. This shows that an early detection of “Artificial Alzheimer” is possible – the process of reaching a judgement on the input – whether it is right or wrong – is longer when the inputs have been distorted and synapses have been deleted. It is also clear that perfect logos as input (blue curve) give uninteresting results. Both these observations are obvious from Figure 4.

**Fig. 4.**
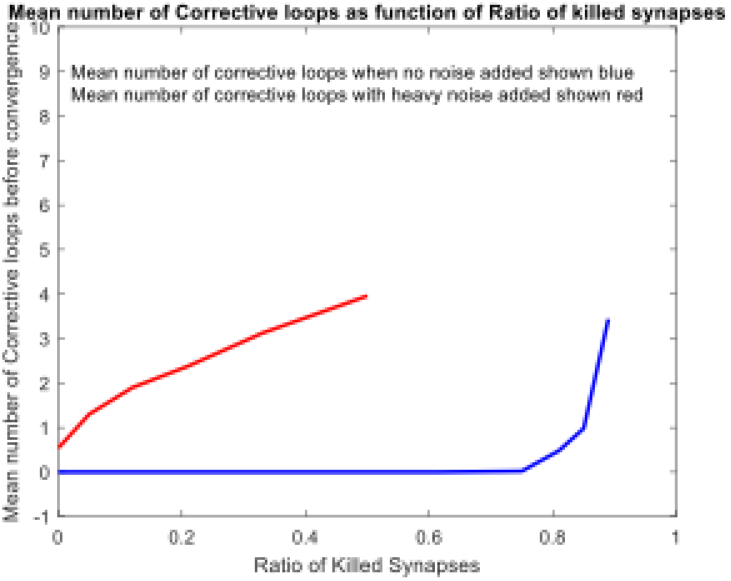
Mean number or corrective loops necessary for convergence when no noise has been added to logos (blue curve) and when heavy noise has been added to logos (red curve). Convergence is not guaranteed for the greatest number of killed neurons (ratio ¿ 0.8).

### 2.3 A surprising – but useless – result

Figure 5 shows that it is perhaps surprising that the quality of the restoration of corrupted logos deteriorate much “later” i.e. with more deletions of synapses. It is not surprising that perfect identifications is better when no noise is added (blue curve) than when heavy noise is added to the inputs (red curve). It is surprising, however, that when heavy noise is added, slightly better results are reached when some synapses have been killed.

**Fig. 5.**
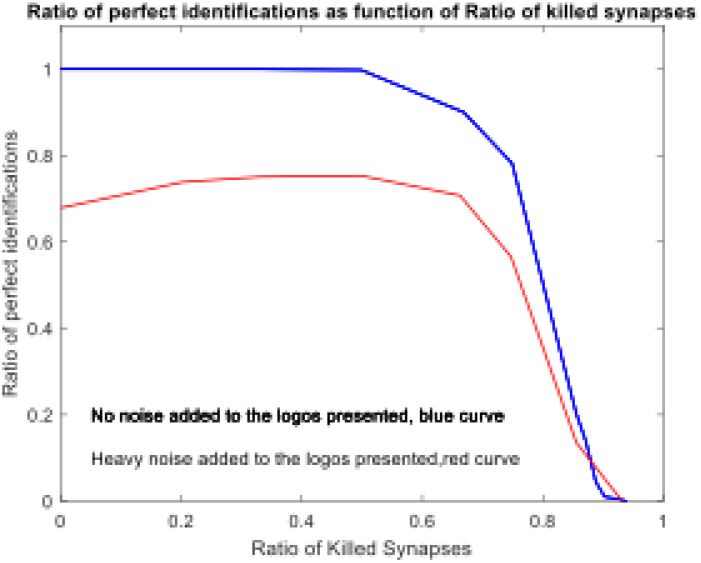
Ratio of perfect identifications as function of killed neurons when no noise was added to logos (blue curve) and when heavy noise was added to logos (red curve).

## 3 Conclusions

### A Suggestion for diagnosing the real thing – Alzheimer’s disease

What would be needed? A set of logos that most people have learned well. The set of ten digits or the set of 24 letters in the latin alphabet as used in English would suffice for literate people in the West. Such symbols could be corrupted by noise and presented to test persons. It would conceivably take longer time to read them out than reading out uncorrupted symbols and the increase in time could be a measure of the cognitive decline in test persons. In the test single distorted letters should be shown, not in alphabetical order since the alphabetical order would offer important clues to most test persons. Undistorted and distorted letters are shown in Figure 6. A sample test page is shown in Figure 7.

**Fig. 6.**
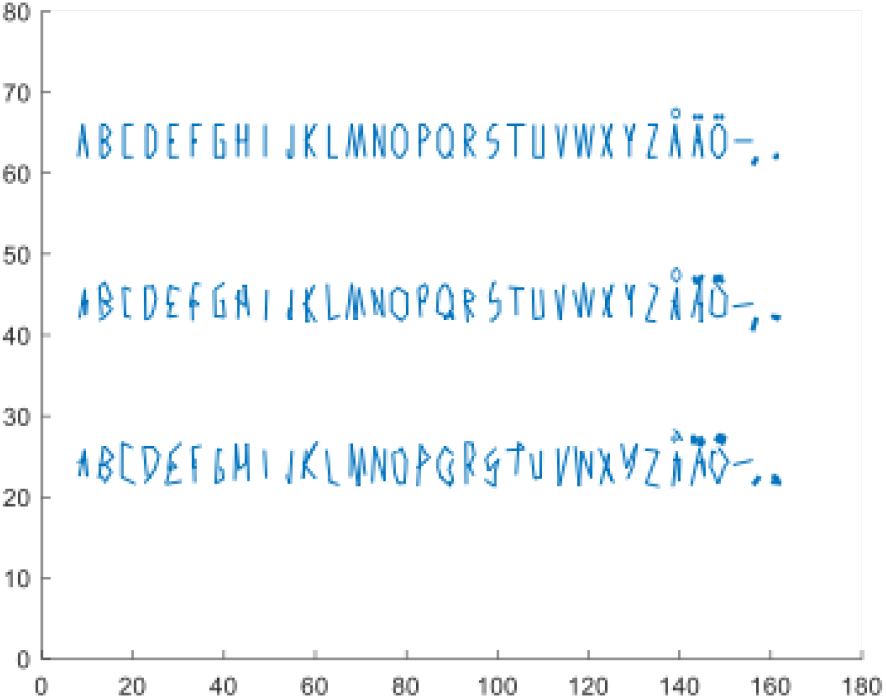
The Swedish alphabet, undistorted and with two degrees of distortion.

**Fig. 7.**
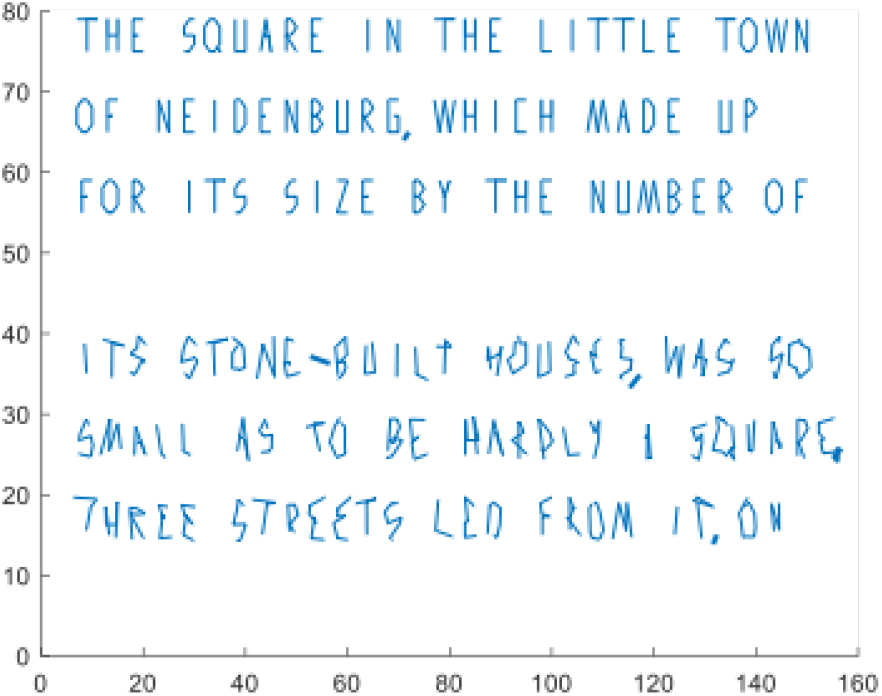
A sample test page.

It is conceivable that reading texts with undistorted letters and texts with distorted letters also could yield interesting results as well. In this case context will give strong clues just as alphabetical order would – it would be a test also of cognition on a “higher level” than merely reading letters.

## References

[1] Horn, D., Ruppin, E., Usher, M., Herrmann, M.: Neural network modeling of memory deterioration in Alzheimer’s disease. Neural computation 5(5), 736–749 (1993)

[2] Duch, W.: Therapeutic applications of computer models of brain activity for Alzheimer disease. (2000)

[3] Hopfield, J.J.: Neural networks and physical systems with emergent collective computational abilities. Proceedings of the national academy of sciences 79(8), 2554–2558 (1982)

[4] Haykin, S., Network, N.: A comprehensive foundation. Neural networks 2(2004), 41 (2004)

